# Insertion sequences drive the emergence of a highly adapted human pathogen

**DOI:** 10.1101/452334

**Authors:** Erwin Sentausa, Pauline Basso, Alice Berry, Annie Adrait, Gwendoline Bellement, Yohann Couté, Stephen Lory, Sylvie Elsen, Ina Attrée

## Abstract

Taxonomic outliers of *Pseudomonas aeruginosa* of environmental origin have recently emerged as infectious for humans. Here we present the first genome-wide analysis of an isolate that caused fatal hemorrhagic pneumonia. We demonstrate that, in two sequential clones, CLJ1 and CLJ3, recovered from a patient with chronic pulmonary disease, insertion of a mobile genetic element into the *P. aeruginosa* chromosome affected major virulence-associated phenotypes and led to increased resistance to antibiotics used to treat the patient. Comparative proteome and transcriptome analyses revealed that this insertion sequence, ISL3, disrupted genes encoding flagellar components, type IV pili, O-specific antigens, translesion polymerase and enzymes producing hydrogen cyanide. CLJ3 possessed seven fold more IS insertions than CLJ1, some modifying its susceptibility to antibiotics by disrupting the genes for the outer-membrane porin OprD and the regulator of β-lactamase expression AmpD. In the *Galleria mellonella* larvae model, the two strains displayed different levels of virulence, with CLJ1 being highly pathogenic. This work reveals ISs as major players in enhancing the pathogenic potential of a *P. aeruginosa* taxonomic outlier by modulating both, the virulence and the resistance to antimicrobials, and explains the ability of this bacterium to adapt from the environment to a human host.

## Introduction

Emerging infectious diseases caused by multidrug resistant bacteria represent a serious threat for human well-being and health. Several hundred novel pathologies caused by infectious agents have been reported during the last forty years (1). Environmental bacteria can adapt to a human host by acquisition of virulence determinants through chromosomal rearrangements due to mobile genetic elements, horizontal gene transfer, or small local sequence changes including single nucleotide substitutions, or horizontal gene transfer (2, 3).

The *Pseudomonas* genus is one of the most important groups of bacteria thriving in diverse environments capable of causing plant or animal diseases. Species such as *P. aeruginosa, Pseudomonas fluorescens*, and *Pseudomonas syringae* adversely impact human health and agriculture (4–6). *P. aeruginosa* is a particularly successful opportunistic pathogen found frequently in humid environments associated with human activities. In hospital settings, infections caused by multi-resistant *P. aeruginosa* strains present a real danger for elderly individuals, patients undergoing immunosuppressive therapies and those requiring treatment with invasive devices in Intensive Care Units. In addition to acute infections, *P. aeruginosa* is a common cause of chronic wound infections as well as long lasting respiratory infections of patients with cystic fibrosis (CF) and chronic obstructive pulmonary disease (COPD). During chronic infections the bacteria adapt to the particular host environment by changing metabolic pathways and synthesis of virulence-associated components (7). Additionally, in-patient evolution involves acquisition of loss-of-function mutations in genes of motility, antibiotic resistance, acute virulence and envelope biogenesis (8–12).

Recent massive whole genome sequencing allowed classification of clinical and environmental *P. aeruginosa* strains in three clades (13, 14). The pathogenic strategies of the three clades rely on different toxins. The two most populated clades inject the toxins, also referred to as effectors ExoS, ExoT, ExoY, and ExoU, directly into host cell cytoplasm through a complex molecular syringe using type III secretion system (T3SS) (15). The third clade is occupied by taxonomic outliers that lack all the genes encoding the effectors and the components of the T3SS. The first fully sequenced taxonomic outlier, PA7, was multi-drug resistant and non-virulent in an acute lung infection mouse model (16, 17). Other PA7-related strains were mainly of environmental origin (18, 19) or associated with both, acute (wounds and urinary tract) and chronic (CF and COPD) human infections (20–22). They recently emerged as highly virulent for humans potentially through the secretion of a pore-forming toxin Exolysin, ExlA (16, 21–23). The most pathogenic *exlA*^+^ *P. aeruginosa* strain described up to date is the strain CLJ1 isolated at the University Hospital in Grenoble, France, from a COPD patient suffering from hemorrhagic pneumonia (16). In murine acute lung infection model, CLJ1-infected lungs featured an extensive damage to endothelial monolayers, bacteria transmigrated into the blood and disseminated into secondary organs without being detected by the immune system. This differs greatly from consequences observed by the T3SS^+^ strain PAO1 (16, 24).

To get insights into molecular determinants of pathogenesis expressed by the Exolysinproducing *P. aeruginosa* taxonomic outliers and to assess the extent of evolutionary adaptation during the course of infection, we performed a comprehensive comparative genome-wide study of two clonal variants, CLJ1 and CLJ3, isolated from the same patient at different time-points during hospitalization. The gathered data demonstrated that mobile genetic elements belonging to the ISL3 family insertion sequences (IS), originally found in soil bacteria *Pseudomonas stutzeri* and *Pseudomonas putida*, shape the virulence traits and strategies employed by those strains to colonize and to adapt to the human host.

## Materials and methods

### Bacterial strains and culture conditions

*P. aeruginosa* strains used in this study are CLJ1 and CLJ3 (16). Bacteria were grown at 37°C in liquid Lysogeny Broth (LB) medium (10 g/L Bacto tryptone, 5 g/L yeast extract, 10 g/L NaCl) with agitation until the cultures reached an optical density at 600 nm (OD_600_) of 1.0 unless indicated. For the assessment of c-di-GMP levels, pUCP22-p*cdrA-gfp*(ASV)^c^ was introduced into CLJ1 and CLJ3 strains by electroporation, as previously described (25), and selected on LB agar plates containing 200 μg/mL of carbenicillin.

### Genome analysis

The details on sequencing, assembly, annotation and comparison are described in Supplementary materials and methods. The assembly and annotation statistics for the CLJ1 and CLJ3 genomes are presented in Table S1. Functional annotation was performed on the Rapid Annotations based on Subsystem Technology (RAST) Genome Annotation Server version 2.0 (26) using Classic RAST annotation scheme and GLIMMER-3 gene caller. Manual curation was done based on the annotations of orthologous genes in PA7 and PAO1 strains from the Pseudomonas Genome Database (27). Circos 0.69-3 (28) was used to create multiomic data visualizations in Fig. 1 and Fig. S1. The whole genome shotgun projects of CLJ1 and CLJ3 have been deposited at DDBJ/ENA/GenBank under the accessions PVXJ00000000 and PZJI00000000, respectively.

**Fig. 1.**
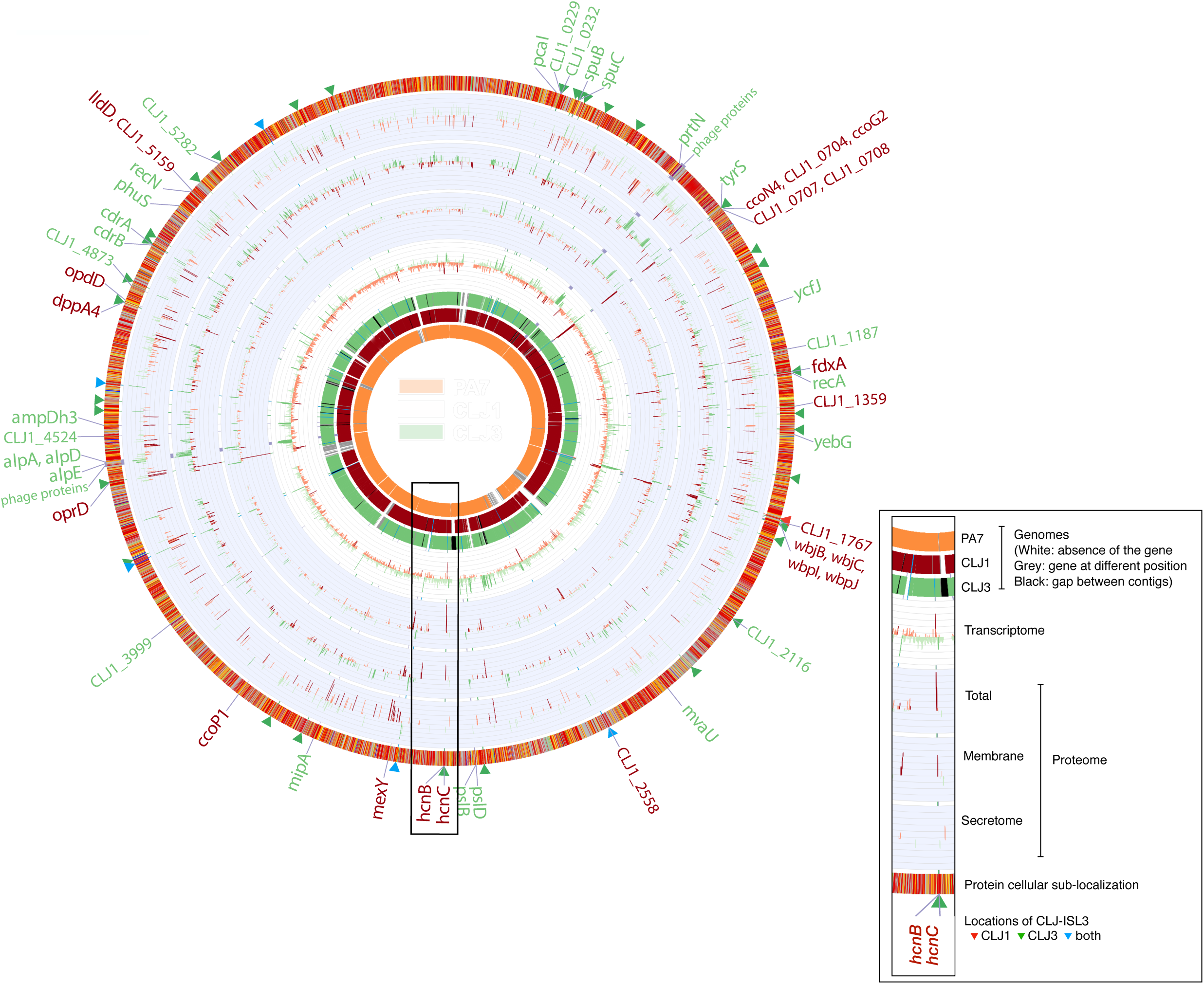
Comparison of the CLJ1 and CLJ3 genome, transcriptome and proteome. The overview of the three genomes, including the reference genome of the PA7 strain, is shown on the left, while the image on the right is an enlarged genomic segment at the hcn locus with a more detailed description of the data. The red bar charts indicate that the gene or protein is more expressed in CLJ1, whereas the green bars show higher expression in CLJ3; darker tone indicates statistically significant expression difference between the two strains (False Discovery Rate < 0.05). The labels linked to the outmost ring show the genes that are differentially expressed in both, RNA-Seq and at least one of the proteomic datasets. The protein subcellular localization (outmost ring) is colored as in the Pseudomonas Genome Database (30). The CLJ-ISL3 insertions are indicated by red, green and blue triangles, depending on their presence in CLJ1, CLJ3 or both strains, respectively.

### Identification of insertion sites for IS elements

CLJ-ISL3 insertion locations in the gaps between CLJ3 contigs were detected by checking for the inverted repeat sequences at the contigs’ ends, while accounting for the shared gene synteny between CLJ1, PA7 and PAO1 genomes. We also used panISa version 0.1.0 (https://github.com/bvalot/panISa) with default parameters to search for ISs in CLJ3 reads, mapped using BWA-MEM algorithm from BWA version 0.7.15 (29) to PA7 and CLJ1 genomes, respectively.

### Transcriptome

The RNA for RNA-Seq was prepared as described (30) from bacterial cultures grown in duplicates in LB to OD_600_ of 1. The preparations of the Illumina libraries and sequencing were done by standard procedures at the Biopolymer Facility, Harvard Medical School, Boston, USA. The analysis was done as described in Supplementary materials and methods.

### Mass spectrometry-based quantitative proteomic analyses

Samples for proteomics were prepared and analysed by nanoliquid chromatography coupled to tandem mass spectrometry (Ultimate 3000 coupled to LTQ-Orbitrap Velos Pro, Thermo Scientific) as described previously (31) with slight modifications, as described in Supplementary materials and methods. Each fraction was controlled by western blot, using appropriate antibodies. The protein content in total, membrane, and secretome proteomes of CLJ1 and CLJ3 were analyzed independently from the others. Statistical analyses were performed using ProStaR (32). In total, a list of 2 852 quantified proteins was obtained. The mass spectrometry proteomics data have been deposited to the ProteomeXchange Consortium via the PRIDE (33) partner repository with the dataset identifier PXD011105.

### Phenotypic analyses

HCN production was assessed on induction plate containing arginine (HCN precursor) as previously described (22). To monitor the c-di-GMP levels, the fluorescence-based reporter plasmid pUCP22-p*CdrA-gfp*(ASV)^c^ was used (34). Bacteria carrying the plasmid were subcultured at OD_600nm_ of 0.05 in black 96-well plate with clear bottom, and incubated at 37°C and 60 rpm in the Fluoroskan reader. The fluorescent emission was measured every 15 min at 527 nm following an excitation at 485 nm for 6 h. Serum sensitivity was assessed by a protocol adapted from (35). Two different human sera from the Etablissement Français du Sang (EFS) were used in all experiments. Overnight cultures of CLJ1 and CLJ3 bacteria were pelleted at 3,000 rpm for 5 min and suspended in Hanks Balanced Salt Solution (HBSS, GIBCO) with 0.1% of gelatin and adjusted to 10^8^ colony forming units (CFU) per mL. The bacteria (10^6^) were incubated in presence of 10% of human serum in a final volume of 3 mL for 15 min or 30 min at 37°C under gentle agitation. A control with heat inactivated serum at 56°C for 30 min was done. CFU were determined at 0, 15, and 30 min by serial dilutions and spreading on LB plates.

### Infections of *Galleria mellonella* larvae

The calibrated wax moth larvae *Galleria mellonella* were purchased from the French company Sud-Est Appats(http://www.sudestappats.fr). Healthy, uniformly white larvae, measuring around 3 cm were selected for infection. The bacteria were grown until the OD_600_ of 1 and diluted in PBS to approximately 10^3^ bacteria/mL. Insulin cartridges were sterilized and filled with bacterial solutions. The larvae were injected with 10 μL of bacterial suspensions using the insulin pen. The exact number of bacteria used in pricking was obtained by spotting five times 10 μL with the pen on agar plates and by counting colony-forming units after growth at 37°C. The infected animals were placed in petri dishes and set at 37°C. The dead larvae were counted over indicated period. Twenty larvae were used per condition and the experiment was performed twice.

### RT-qPCR

To quantify selected transcripts, total RNA from 2.0 mL of cultures (OD_600_ of 1.0) was extracted with the TRIzol Plus RNA Purification Kit (Invitrogen) then treated with DNase I (Amplification Grade, Invitrogen). RT-qPCR was performed as described (36) with few modifications as described in Supplementary materials and methods. The sequences of primers were designed using Primer3Plus (http://www.bioinformatics.nl/cgibin/primer3plus/primer3plus.cgi) and are given in Table S2.

## Results

### Analysis of the CLJ1 genome and its regions of genomic plasticity

Strain CLJ1, an antibiotic-sensitive *P. aeruginosa,* was isolated from the patient with necrotizing hemorrhagic pneumonia. Twelve days later, after initiating antibiotic therapy the patient condition worsened and at this time CLJ3, a multidrug resistant clonal variant, was isolated (16). We have also shown that CLJ1 is a cytotoxic strain that shares the main genomic features with the first fully sequenced antibiotic resistant taxonomic outlier PA7 (16, 17). Notably, CLJ1 lacks the entire locus encoding the T3SS machinery and the genes encoding all known T3SS effectors; however, it carries the determinant for the two-partner secretion pore-forming toxin, Exolysin. To initiate genome-wide studies on mechanisms conferring the specific phenotypes of CLJ1, we fully sequenced the genome of this strain and compared it to PA7. As expected, most of the core genes of CLJ1 are similar to PA7 (Fig. 1 and Fig. S1). However, the content and the distribution of several regions of genomic plasticity (RGPs) (37) are different. CLJ1 genome contains 15 regions that are absent from the PA7 genome and lacks 26 PA7-specific regions. Among differences between CLJ1 and PA7 genomes there is a putative integrated plasmid carrying several genes that confers aminoglycoside resistance, regions encoding two type I restriction-modification systems, a pyocin protein, and a mercury resistance system (Tables S3 and S4). All CLJ1 specific regions (CLJ-SR), except CLJ-SR14, are present in other *P. aeruginosa* strains, some being phylogenetically unrelated to the PA7 strain. Strikingly, CLJ-SR14 carries 55 genes, many predicted to encode proteins involved in metabolism and resistance to heavy metals (Table S5), suggesting environmental origin of the strain. Modifications in three replacement islands (38, 39) can mediate the CLJ1-specific phenotype. The CLJ replacement island in RGP60 (harboring pilin and pilin modification genes) carries a group I pilin allele, unlike PA7 that has a group IV allele (40). CLJ1 encodes, within RGP9, a b-type flagellin as the principal component of its flagellum, while PA7 possesses a-type flagellin (17). Both, RGP9 containing the flagellin glycosylation genes and the replacement island in RGP31 bearing the O-specific antigen (OSA) biosynthesis gene cluster are further modified in CLJ strains (see below). While most of the genes within RGP7 (pKLC102-like island) are missing in CLJ1, the so-called Dit island previously found in a CF isolate of *P. aeruginosa* (37) is inserted in 5’ region of RGP27 at the tRNA^Gly^. The determinants encoded in this island provide the bacteria the ability to degrade aromatic diterpenes, which are tricyclic resin acids produced by wounded trees, and to use them as sole carbon and energy source (41). The Dit island is uncommon in the genomes of *P. aeruginosa* strains, but it is frequently present in different soil bacteria such as *Burkholderia xenovorans*, *P. fluorescens*, and *Pseudomonas mendocina*, further implying the environmental origin of the CLJ1 strain. Among potential virulence-associated genes, CLJ1 lacks one of the type 6 secretion system loci (*PSPA7_2884-2917*) encoding an injection machine for toxins active against both, prokaryotes and eukaryotic cells (42). The *plcH* and *plcR* genes, coding for the hemolytic phospholipase C precursor and its accessory protein respectively, are also absent from the CLJ1 genome. Whereas *cupA* fimbrial gene cluster (*CLJ1_2899-2903*; *PSPA7_3019-3023*) is present in the remodeled RGP23, the entire *cupE1-6* operon (*PSPA7_5297-5302*) encoding cell surface fimbriae involved in the maintenance of a biofilm structure (43) is missing.

### Evidence of mobile genetic elements in the key pathogenic regions

Following infection, CLJ1 strain provokes striking damage to mouse lung tissues without an immunological response by the host (24). The sequential isolate CLJ3 was sampled from the same patient after a series of aggressive antibiotic treatments and displayed different cytotoxicity and antibiotic-sensitivity profiles compared to CLJ1 (16). The genomes of these two strains are almost identical, with a reciprocal best-hit average nucleotide identity (44) between them estimated to be 99.97% (Fig. 1). When analyzing the genomes of CLJ1 and CLJ3, the most striking feature is the presence of multiple copies of a 2,985-bp fragment corresponding to a mobile genetic element, absent from the reference PA7 genome. This sequence is 99% identical to ISL3 family of insertion sequences ISPst2 of *Pseudomonas stutzeri*, ISpu12 of *Pseudomonas putida* and IS1396 of *Serratia marcescens* found in the ISfinder database (45). However, besides coding for a transposase, this CLJ-ISL3 encodes a putative inner membrane protein with seven transmembrane helices from the permease superfamily cl17795 (with the conserved domain COG0701 (46)) and a putative transcriptional metalloregulator of the ArsR family (Fig. 2A). The element is flanked by a pair of 24-bp imperfect inverted repeat sequences GGGTATCCGGAATTTCTGGTTGAT (left inverted repeat, IRL) and GGGTATACGGATTTAATGGTTGAT (right inverted repeat, IRR). Examination of publicly available genome sequences showed that the complete CLJ-ISL3 is also present in a few other bacteria, including a multi-drug resistant *Acinetobacter baumannii* isolate recovered from bronchoalveolar lavage fluid (47).

**Fig. 2.**
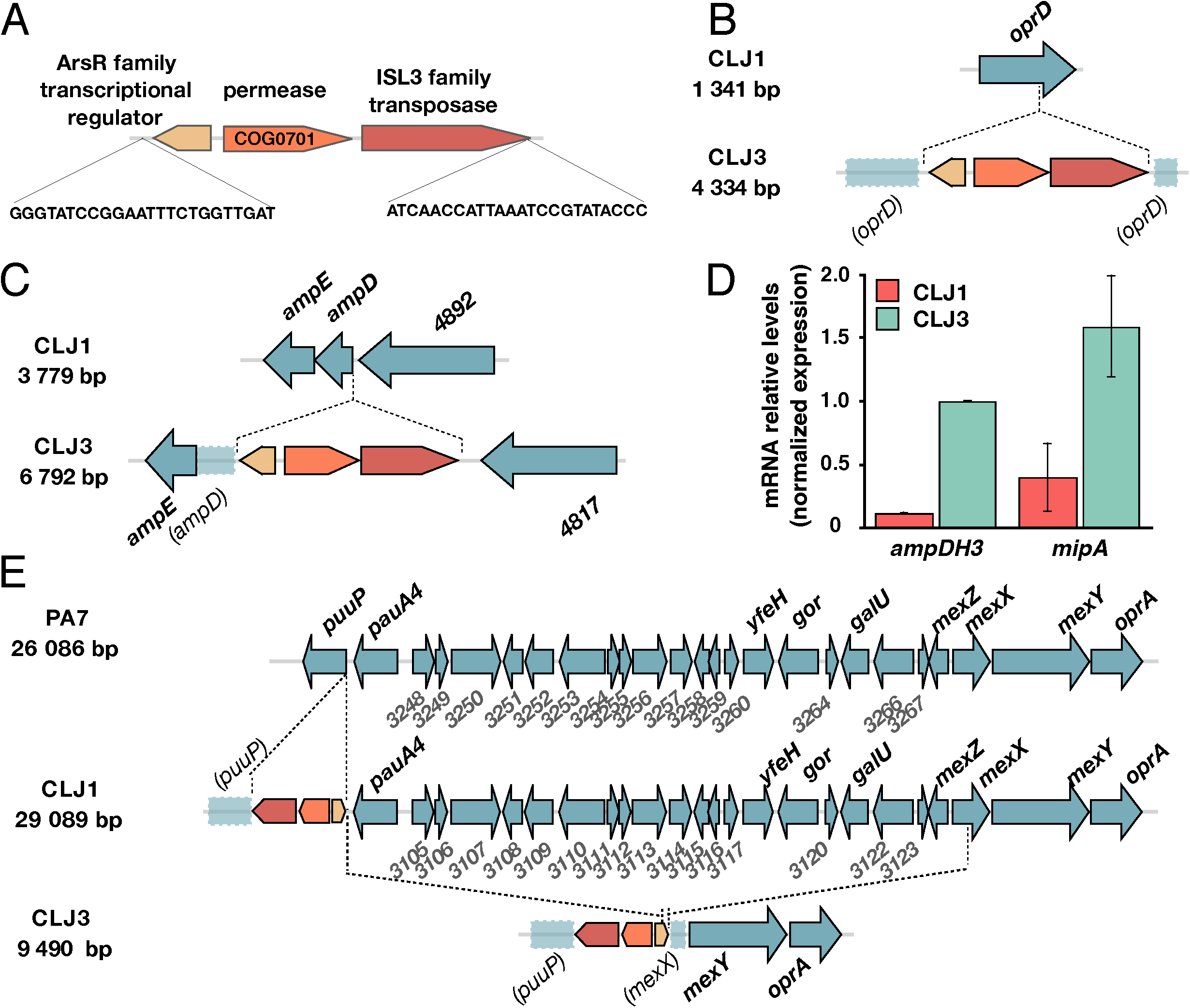
Insertions of CLJ-ISL3 into genes encoding determinants of antibiotic susceptibility. **A** Representation of the 2,985-bp CLJ-ISL3 IS with three genes and inverted repeats at its ends. **B** Representation of the *oprD* disruption in CLJ3 by the CLJ-ISL3 insertion sequence. **C** Location of CLJ-ISL3 in *ampD* in CLJ3. **D** Analysis of the relative expression of *ampDH3* and *mipA* in CLJ1 and CLJ3 strains by RT-qPCR. The bars indicate the standard error of the mean. **E** Gene organization of the region in PA7 and CLJ1 that is absent in CLJ3. CLJ-ISL3 is found in *puuP* in CLJ1 while, both *puuP* and *mexX* are disrupted in CLJ3, with the connecting sequence replaced by CLJ-ISL3 (see the text).

Altogether, we found CLJ-ISL3 in six and forty copies in the genomes of CLJ1 and CLJ3, respectively. The presence of ISs in CLJ3 genome was predicted using a combination of bioinformatics tools, analysis of genome syntheny and detection of inverted sequence (see Materials and Methods). In agreement with the clonal origin of the strains, all CLJISL3 of CLJ1, but one, were located in the same location in CLJ3 genome (Fig. 1 and Table S6). Mass spectrometry (MS)-based quantitative proteomic analyses revealed a strong increase in the level of the transposase protein in CLJ3 compared to CLJ1, that might explain the IS expansion within its genome (Table S7). As mobile genetic elements greatly contribute to phenotypic modifications by changing the gene expression or by gene inactivation (49, 50), we examined their impact on global gene expression and particular phenotypes. To that aim, we compared CLJ1 and CLJ3 transcriptomes and proteomes using respectively RNA-Seq and MS-based quantitative proteomics. The proteomes of three different bacterial fractions (whole bacteria, total membranes, and secretomes) were analyzed. Stringent statistical analyses of extracted data revealed 77 differentially expressed genes/proteins between CLJ1 and CLJ3, both at levels of mRNA and protein (Fig. 1, Tables S7, S8 and S9). Out of these, 27 (35%) are of phage origin, while 32 (42%) are predicted to be localized in bacterial membranes, periplasm or secreted.

### Contribution of CLJ-ISL3 to antibiotic resistance of the CLJ3 strain

The CLJ1 and CLJ3 strains are recent isolates obtained from the patient treated with high doses of different antibiotics without successfully eliminating the infection. CLJ1, isolated before the beginning of the antibiotic therapy, was sensitive to the tested antibiotics, while CLJ3 displayed resistance to most of the antibiotics administrated to the patient (16). To gain insight into mechanisms of antibiotics resistance of CLJ3, we examined the genomic data for gene modifications that could explain the switch in phenotypes to selected antibiotics. We observed that several genes encoding proteins potentially conferring antibiotic resistance are modified by ISs.

As the patient was given different antibiotics of the β-lactam family (ticarcillin, carbapenem, cephalosporin…) and the CLJ3 isolate developed the resistance to all of them, we examined the status of the outer membrane porin OprD (CLJ1_4366) and the chromosomal-encoded AmpC b-lactamase (CLJ1_0728), the two major determinants of intrinsic resistance to this group of antibiotics. We found that *oprD* gene was interrupted by the CLJ-ISL3 (Fig. 2B) resulting in absence of the protein (Table S7). Loss-of-function mutations or deletions in *oprD* are commonly seen in isolates from patients undergoing treatment with imipenem or meropenem (51–54) making the cell envelope impermeable to these antibiotics. Additionally, the identical ISL3 element was found in the 5’ portion of the *ampD* gene (Fig. 2C), encoding the recycling amidase responsible for the production of muropeptide regulators of *ampC* expression (55, 56). Consequently, although higher expression of *ampC* gene was not detected in RNA-Seq, the CLJ3 proteomes contained significantly higher amounts of the AmpC protein compared to those of CLJ1 (Table S7).

Two additional periplasmic proteins involved in peptidoglycan recycling and biosynthesis, the AmpDH3 amidase (CLJ1_5671) (57) and the lytic transglycosylase MltA-interacting protein MipA (CLJ1_3357) (58), were overrepresented in the CLJ3 proteome. In agreement, our datasets also pointed to the overexpression of corresponding genes, further confirmed by RT-qPCR (Fig. 2D). The overproduction of these proteins in a clinical strain suggests their role in adaptation of the strain to host through modulation of peptidoglycan synthesis. However, the molecular mechanisms involved in their increased expression and the significance of their overexpression in bacterial persistence in the host need to be determined.

All *P. aeruginosa* strains carry genes for multiple efflux pumps. The CLJ1/PA7 clade possesses the locus encoding the MexXY-OprA efflux pump which is able to transport multiple antibiotics including fluoroquinolones, aminoglycosides, and certain cephalosporins (reviewed in (59)). However, when compared to CLJ1, there is a *ca*. 20kb deletion in the CLJ3 genome (with a loss of 22 genes corresponding to *PSPA7_3247-3268*) eliminating the entire transcriptional repressor *mexZ* gene and truncating the *mexX* (Fig. 2E). The deleted region is replaced by a copy of CLJ-ISL3 that is bordered by truncated *puuP* and *mexX*. One plausible explanation for the observed genomic arrangement of this region in CLJ3 is that the strain was derived from a yet un-identified clonal strain that had another copy of ISL3 in *mexX*. A recombination between these two IS elements resulted in the 20 kb deletion, simultaneously eliminating the repressor and a portion of *mexX* genes, making this efflux pump nonfunctional. The truncation of the *puuP* gene encoding putrescine importer/permease indicates that the CLJ-ISL3 sequence affects transport of polyamine, which have multiple roles in pathogen biology, including resistance to some antibacterial agents (60).

### Modifications of the O-specific antigens (OSA) cluster due to ISs

OSA is a component of lipopolysaccharide (LPS) and an integral component of the *P. aeruginosa* cell envelope. The OSA biosynthetic gene cluster of CLJ in RGP31 is similar in content to that of PA7, consequently both PA7 and CLJ1 belong to serotype O12 (22). However, insertion of two different ISs alters the gene content and expression levels of the encoded proteins (Fig. 3A). In CLJ1, the ISL3 element was found within the gene encoding NAD-dependent epimerase/dehydratase (CLJ1_1762 corresponding to PSPA7_1970), whereas in the corresponding CLJ3 locus two copies were present, one disrupting *wbjL* and the second located in the intergenic region between *CLJ3_1919* and *CLJ3_1920* (*PSPA7_1972*). Moreover, in CLJ3, two copies of another IS from the IS66 family were also found within the gene *wbjB (CLJ1_1771; CLJ3_1930)* and downstream of *wbjM (CLJ1_1778; CLJ3_1936)*. These insertions negatively affect expression of these genes, or mRNA stability, as their mRNA and protein levels were higher in CLJ1 than in CLJ3 (Tables S7 and S8).

**Fig. 3.**
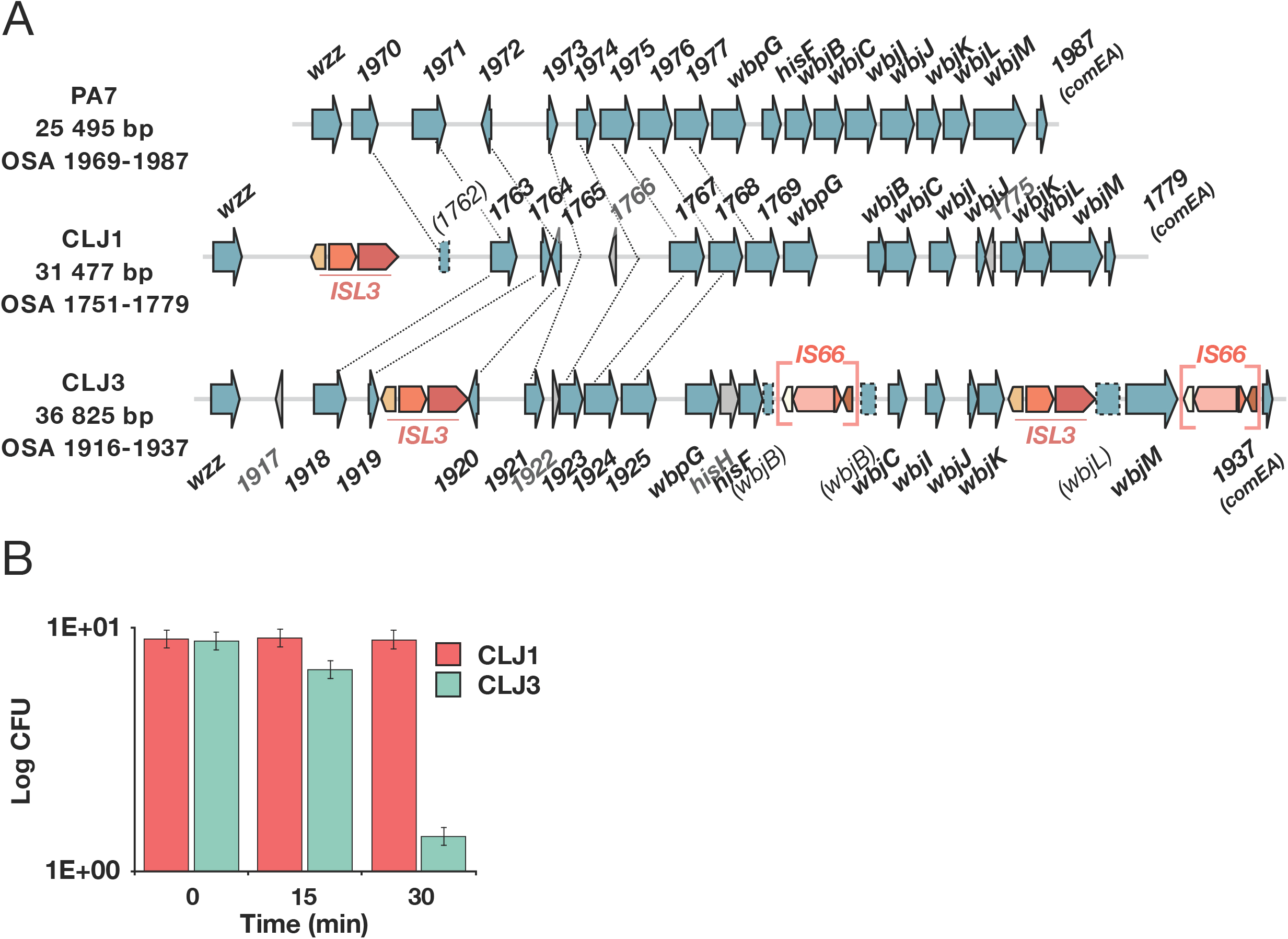
ISs within OSA loci and impact on serum sensitivity. **A** Comparison of the OSA region in PA7 and CLJ strains. CLJ-ISL3 is inserted in one location in CLJ1 and at two sites in CLJ3, while IS66 is present in two copies in CLJ3. Correspondence between non-annotated genes is depicted with connecting lines. **B** Kinetics of serum killing of CLJ1 and CLJ3 strains. Bacteria were incubated with human sera for indicated time, diluted and spotted on agar plates. Colony forming units were counted after incubation at 37°C.

Furthermore, some other proteins encoded in the 3’ part of the OSA cluster, i.e. CLJ1_1767 (PSPA7_1975), WbjB (CLJ1_1771), WbjC (CLJ1_1772), WbjI (CLJ1_1773) and WbjJ (CLJ1_1774) were more abundant in CLJ1 (Table S7). Analogous variations of the OSA loci leading to variability of O-antigens have been reported (61, 62). Interestingly, the previously described deletion at the *mexX*-*puuP* locus in CLJ3 encompasses also the *galU* gene (*CLJ1_3121*, Fig. 2E), encoding UDP glucose pyrophosphorylase (63) that adds sugar moieties onto the inner core on lipid A, serving as the anchor for the full length LPS. The LPS truncation due to absence of *galU* leads to a so-called rough LPS, higher susceptibility to serum-mediated killing and reduced *in vivo* virulence (64). The CLJ3 strain was non-agglutinable and sensitive to serum compared to the CLJ1 strain (Fig. 3B). The *in vivo* selection for acquisition of a serum sensitivity phenotype is unclear but it may reflect the adaptation of CLJ3 to the host environment through resistance to phages and antibiotic challenges, as it has been reported for some other clinical strains (65). Modification in LPS structures in *P. aeruginosa* strains chronically infecting CF patients is considered as an adaptation mechanism to a more “persistent” lifestyle, with the LPS molecule being less inflammatory (64). Thus, multiple mechanisms contributed to final LPS structure in CLJ clones. Those events could have provided to strains variable advantages during infection process such as resistance to antimicrobials or altered recognition by the immune system.

### ISs determine the repertoire of surface appendices

Inspection of CLJ1 and CLJ3 genome sequences showed that CLJ-ISL3 has strongly affected the flagellar biosynthetic locus in both strains. The CLJ3 genome revealed an organization and gene content of the flagellar locus similar to that found in PAO1, which synthesizes a b-type flagellum (66), but with two copies of CLJ-ISL3 (Fig. 4A). The first IS interrupts *flgL (CLJ1_4222)*, encoding the flagellar hook-associated protein that enables the anchoring of the flagellum to the cell envelope (67), while the second IS element is in *fgtA (CLJ1_4219)* coding for the flagellin glycosyl transferase. The two ISs seem to have recombined in CLJ1 creating a deletion between truncated *fgtA* and *flgL* genes (Fig. 4A). This recombination event suggests that CLJ1 is not the direct ancestor of CLJ3, but that the two strains evolved from a common ancestor. The absence of flagellum is in agreement with the non-motile phenotype of the strain observed during eukaryotic cell infection and on soft agar (22). Bacterial flagella also modulate immune response during the infection, because the major flagellar subunit flagellin binds to TLR5 and initiates the TLR-dependent signaling and activation of expression of pro-inflammatory cytokines (68). The absence of assembled flagella in the isolates explains the lack of detectable pro-inflammatory cytokine interleukin-1β (IL-1β) and TNF in bronchoalveolar lavages of CLJ1-infected mice (23, 24). CLJ1 is also devoid of twitching mobility (22); therefore we examined the genomic data for possible mutations in genes encoding type IV pili (T4P) and found an insertion of CLJ-ISL3 within the 5’ part of *pilM* gene in both CLJ1 and CLJ3 genomes (Fig. 4B). This gene, encoding a cytoplasmic actin-like protein, is the first gene of the *pilMNOPQ* operon reported to be essential for both T4P biogenesis and twitching motility (69). Expression of the entire operon, likely due to the polar effect of the IS element on the downstream genes, seems to be affected since no proteins were detected by proteomic analysis, unlike many other Pil proteins encoded in other operons (Table S7). This finding was intriguing as the action of the toxin Exolysin relies greatly on T4P in another *exlA*+ strain, IHMA87 (70). Therefore, we examined CLJ1 proteomes for the presence of other putative adhesive molecules that may substitute for the T4P function during host cell intoxication. Based on proteomic datasets, the CLJ1 strain synthesizes components of five two-partner secretion (TPS) systems whose predicted secreted components are annotated as haemolysins/haemagglutinins, including the Exolysin (CLJ1_4479) responsible for CLJ1 cytotoxicity CdrA and CdrB (CLJ1_4999 and CLJ1_5000) are significantly overrepresented in the CLJ3 strain as revealed by proteomics and RNA-Seq analysis (Tables S7 and S8). In the PAO1 strain, the adhesin CdrA is regulated by the secondary messenger cyclic-di-GMP (c-di-GMP), and its expression is increased in biofilm-growing condition (71). We assessed the cellular levels of c-di-GMP by using a reporter that is based on the transcriptionally fused c-di-GMP-responsive *cdrA* promoter to a gene encoding unstable green fluorescent protein (34). The p*cdrA*-*gfp* (ASV)^C^ monitoring plasmid clearly showed higher levels of the second messenger in CLJ3 during growth (Fig. 4C), suggesting that CLJ3 may have adapted to host conditions by switching to biofilm lifestyle. Another TpsA protein detected in CLJ secretomes is CLJ1_4560, a HMW-like adhesin recently named PdtA in the PAO1 strain. PdtA plays a role in *P. aeruginosa* virulence as demonstrated by using *Caenorhabditis elegans* model of infection (72). The production of five TPSs, including the protease LepA (CLJ1_4911) (73) and at least one contact-dependent inhibition protein Cdi (CLJ1_2745) (74, 75), in CLJ strains (Table S7) indicates that this family of proteins may play an important role during colonization and infection, but their respective contributions in adhesion, cytotoxicity, or inter-bacterial competition during infection process need to be investigated.

**Fig. 4.**
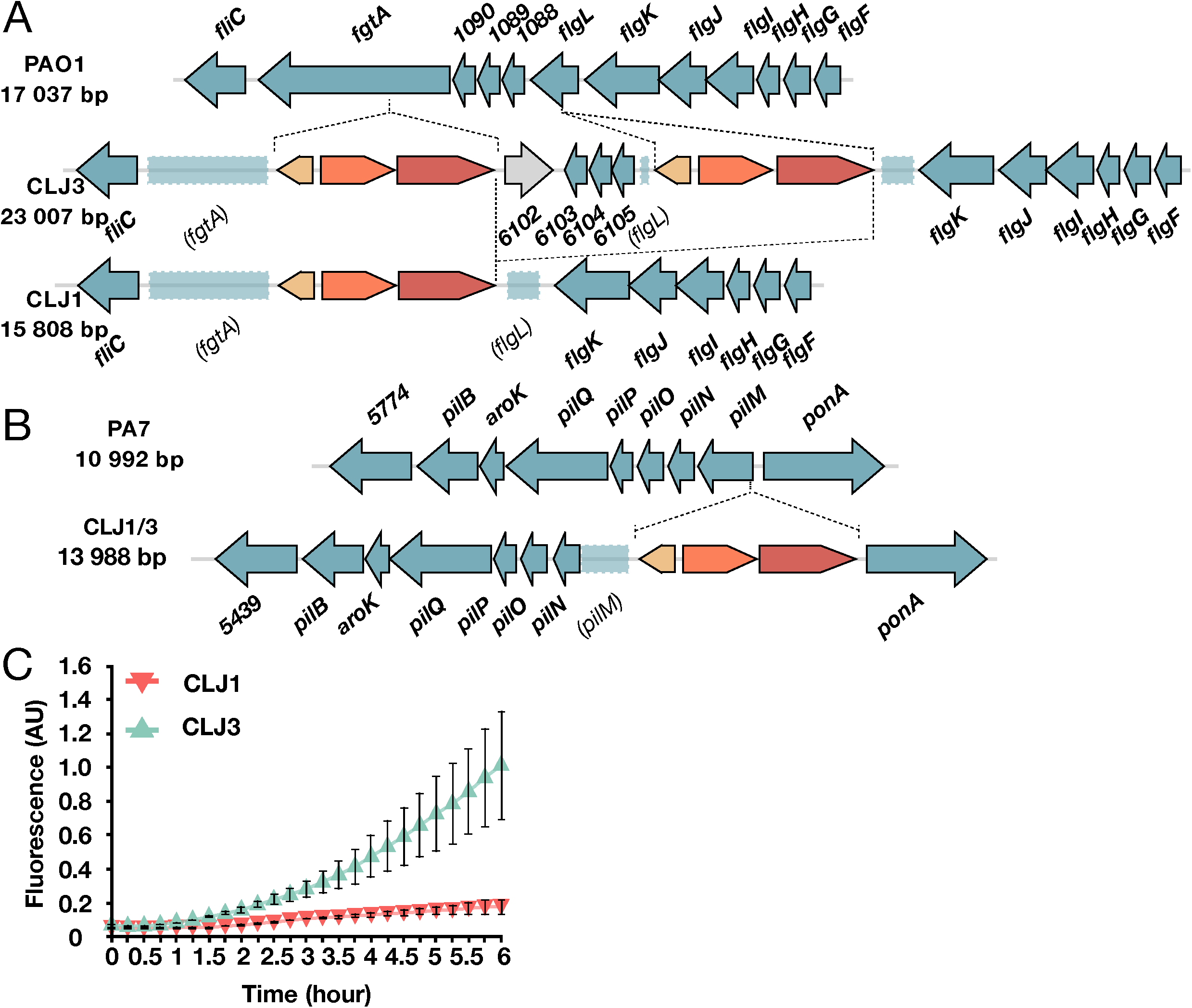
Modifications in components associated with surface appendices. **A** Gene organization of *flgL* region in PAO1 and the CLJ1 strain. Two CLJ-ISL3, interrupting *fgtA* and *flgL*, were found in the CLJ3 genome that probably recombined in CLJ1, leaving only one copy of the IS. **B** Representation of the *pilMNOPQ* operon. The CLJ-ISL3 in *pilM* gene was found in CLJ1 and CLJ3. **C** Synthesis of the adhesin CdrA is linked to high intracellular c-di-GMP levels. The c-di-GMP levels in CLJ1 and CLJ3 strains were monitored using the pcdrA-gfp(ASV)c plasmid. Fluorescence was measured every 15 min for 6 h of growth. The error-bars indicate the standard deviation.

### Other putative virulence factors, phages and metabolism

RNA-Seq results supported by proteomics data showed an increase in expression of the enzymes HcnB (CLJ1_2955) and HcnC (CLJ1_2954) in the CLJ1 strain (Table S9). The inspection of the *hcn* operon revealed that the CLJ-ISL3 element was inserted in the 5’ part of the *hcnB* gene in CLJ3 genome (Fig. 5A), in agreement with absence of expression measured using RT-qPCR (Fig. 5B). The *hcn* genes encode the subunits of hydrogen cyanide (HCN) synthase that produce this toxic secondary metabolite (76). To detect the HCN produced by the strains, a paper impregnated with a reaction mixture containing Cu^2+^ ions was placed above bacteria seeded onto agar plates, a white-to-blue color transition being indicative of HCN in the gas phase. In agreement with proteomic data, CLJ1 was able to produce HCN in higher quantities than CLJ3 and other *P. aeruginosa* strains from the *exlA*^+^ collection (Fig. 5B, (22)). Many *P. aeruginosa* isolates from individuals with CF produce high levels of HCN (77) and the molecule has been detected in the sputum of *P. aeruginosa*-infected CF and bronchiectasis patients (78, 79). HCN is also a regulatory molecule, capable of inducing and repressing other genes (80), including a cluster of genes *PA4129*-*PA4134* in PAO1. This cluster (*CLJ1_0701-CLJ1_0708*) is expressed at higher levels in CLJ1 compared to CLJ3 as revealed by RNA-Seq and quantitative proteomics (Fig. 1, Tables S7 and S8) and confirmed for *ccoG2* and *ccoN4* genes by RT-qPCR (Fig. 5C). The *ccoN4* gene belongs to the *ccoN4Q4* operon, one of the two *ccoNQ* orphan gene clusters present in *P. aeruginosa* genome. Its upregulation in the cyanogenic CLJ1 is in agreement with the study showing that, although *P. aeruginosa* encodes a cyanide-insensitive oxidase CIO, isoforms of *cbb_3_*-type cytochrome *c* oxidase containing CcoN4 subunit were produced in response to cyanide, exhibiting higher tolerance towards this poisonous molecule in low-oxygen conditions (81).

**Fig. 5.**
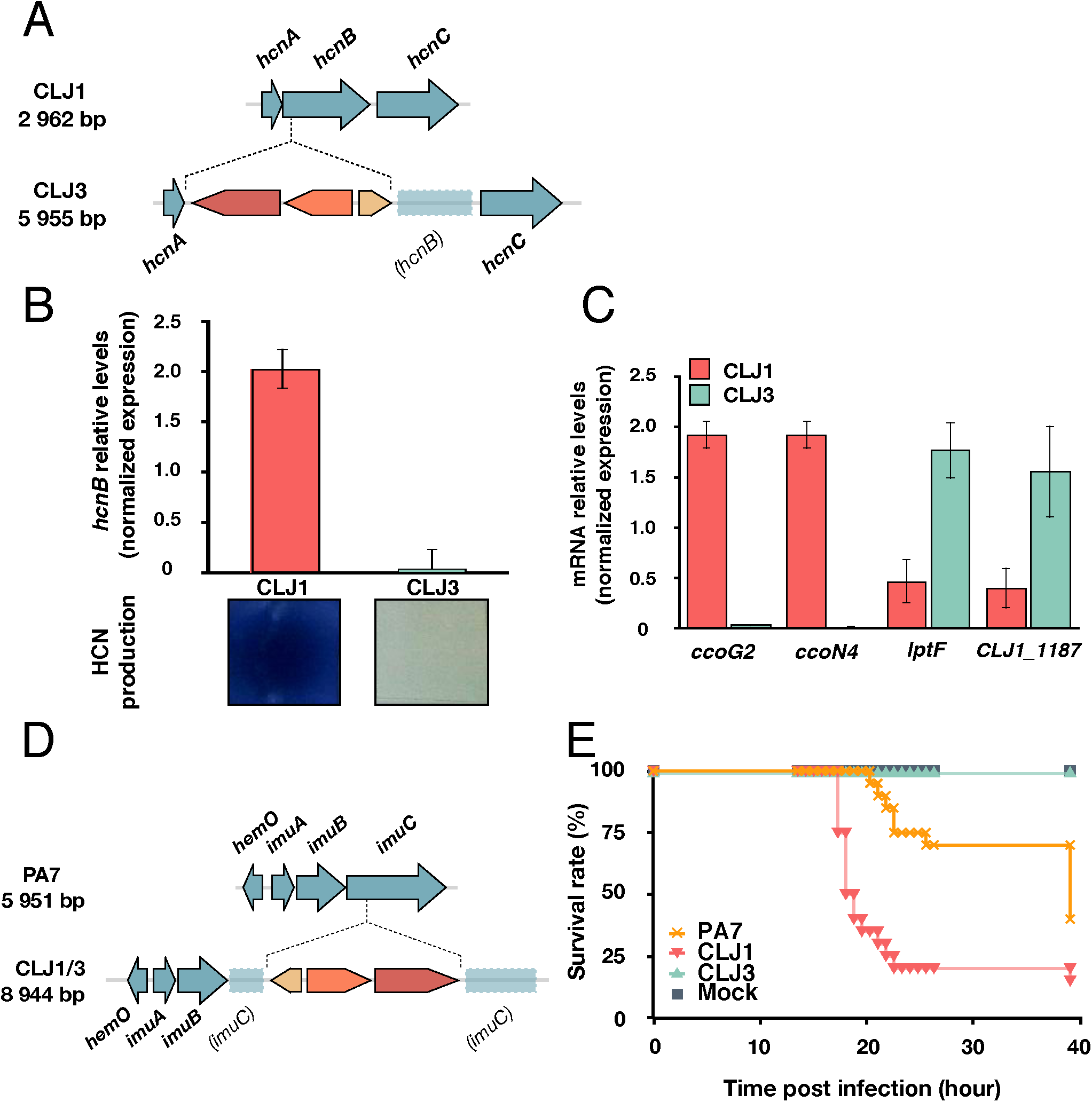
Expression of virulence factors and pathogenicity. **A** The *hcn* operon in CLJ1 and insertion of CLJ-ISL3 within the *hcnB* gene in CLJ3. **B** Relative expression of *hcnB* measured by RT-qPCR and hydrogen cyanide production in indicated strains. **C** Analysis of the relative expression of *ccoG2*, *ccoN4*, *lptF*, and CLJ1_*1187* in CLJ1 and CLJ3 strains by RT-qPCR. The bars indicate the standard error of the mean. **D** The *imuABC* operon and insertion of CLJ-ISL3. **E** Survival of *Galleria* following injection of different strains. Twenty larvae were infected with 5-10 bacteria (estimated from CFU counts) and their survival was followed over indicated period by counting the dead ones. PBS was injected as mock control.

Examining the data for differential expression of putative virulence determinants indicated that CLJ3 overproduces Lipotoxin F (LptF, CLJ1_1186), an outer-membrane protein contributing to adhesion to epithelial A549 cells and known to activate host inflammatory response (82). The *lptF* gene is located in a putative operon together with a gene encoding a hypothetical protein predicted to be a lipoprotein located in the periplasm (CLJ1_1187) that was found upregulated in CLJ3 strain by proteomic analyses (Table S7). As *lptF* upregulation was not detected by initial RNA-Seq, we performed RT-qPCRs that demonstrated significant overexpression of both genes *lptF* and *CLJ1_1187*/*CLJ3_1413* in CLJ3 (Fig. 5C) suggesting that Lipotoxin F together with CLJ1_1187 contributed to bacterial adaptation to in-patient environments which is in agreement with increased *lptF* expression in CF isolates (83).

Finally, prophages play important roles in *P. aeruginosa* physiology, adaptation and virulence (38, 84, 85). In the CLJ genomes, there are at least eight regions related to phages (Table S10) and five of them are different or absent from PA7 (Table S3). Transcriptomic and proteomic approaches revealed that twenty-seven phage-related proteins from *CLJ1_0539-0556* in RGP3 and *CLJ1_4296-4314*, including CLJ-SR11, are significantly more abundant in CLJ3 compared to CLJ1 (Tables S7, S8 and S9). Interestingly, the genes encoding bacteriocins, namely pyocins S2, S4, and S5, are absent from CLJ strains. Nevertheless, the activator of pyocin biosynthetic genes, PrtN (CLJ1_0535) (86), is more highly expressed in CLJ3 than in CLJ1. Indeed, the *prtN* gene is located in a region encoding phage-related proteins which is activated in CLJ3 (Fig. 1, Table S9). In both CLJ genomes, we also identified the genes for AlpR (CLJ1_4295, CLJ3_4269) and AlpA (CLJ1_4296, CLJ3_4268), transcriptional regulators of a programmed cell death pathway in PAO1 (87). Finally, *alpA* and all the genes of the *alpBCDE* lysis cassette are highly expressed in CLJ3, and we noticed high propensity of this strain for lysis (Fig. 1, Table S9).

### CLJ-ISL3 inactivates the *imu* operon encoding translesion synthesis machinery

Another location of the ISL3 element in the CLJ1 and CLJ3 genomes is within the *imu* operon also named the “mutagenesis” cassette (Fig. 5D). The *imu* operon encodes the ImuC polymerase (formerly called DnaE2) and other components of translesion synthesis (TLS), which can bypass lesions caused by DNA damage, and consequently, is mutagenic. The ImuC polymerase in *Pseudomonas* contributes to the tolerance to DNA alkylation agents (88), and the inactivation of the operon could limit accumulation of mutations. Interestingly, CF *P. aeruginosa* isolates frequently display a hyper-mutator phenotype, that is primary due to inactivation of *mutS* and has been previously suggested as being advantageous for bacterial adaptation to the CF lungs (89). The *mutS* gene is identical and intact in CLJ1 and CLJ3 and the predicted proteins differ by one amino acid at position 593 from MutS of PA7 (threonine in PA7, serine in CLJ1/CLJ3). Therefore, the physiological impact of inactivation of the translesion synthesis system is unclear. Two additional DNA-repair proteins, RecN (CLJ1_5150) and RecA (CLJ1_1263), were found overrepresented in CLJ3 at the transcriptome and proteome levels (Table S9), suggesting the interplay between different ways of defense mechanism against uncontrolled mutational rates induced by the hostile environment. We found more than 600 single nucleotide polymorphisms (SNPs) between CLJ1 and CLJ3. This is about six times higher compared to sequential isolates found in the same CF patients over 8.8 years in one study (10) and almost 10 times higher than mutations in a matched isolate pair, collected 7 and 1/2 years apart from a single patient in another study (12). However, another study showed that some strains isolated from non-CF patients could share between 176 and 736 SNPs (9). Although we could not precisely account for the role of SNPs, they may contribute to differences in gene expression between the two isolates that could not be directly attributed to ISs.

### Two CLJ clones show different pathogenic potential in *Galleria mellonella*

To assess the global virulence of the two strains in a whole organism, we used wax moth *Galleria* infection model, and followed the survival of infected larvae following inoculation with different strains. Under the same infection conditions, the CLJ1 strain was found more virulent than the PA7 strain, while CLJ3 was unable to kill *Galleria* larvae (Fig. 5E). This result shows that CLJ3, while gaining resistance to antimicrobials, has lost its virulence potential, in agreement with our previous observation that CLJ3 is sensitive to serum (Figure 3B) and less cytotoxic due to the loss of the ability to secrete Exolysin (16). More than 40 insertion sites of ISL3 have been detected in the genome of CLJ3, most within or upstream of genes encoding hypothetical proteins or putative regulators (Table S6) and some of them may have influenced fitness of the CLJ3 strain in the *Galleria* model of infection. Thus, we found that ISs have greatly contributed to pathogenicity of the CLJ1 strain and to multi-drug resistance of CLJ3. As the coexistence of the two isolates in the patient lungs is possible, we can conclude that ISs have determined the global success of the CLJ lineage in establishing fatal infection.

## Discussion

*P. aeruginosa* strains belonging to the group of taxonomic outliers are abundant in humid environments (18, 19, 90, 91) and are considered to be innocuous, based on a previous study (92). Here we present the results of a multiomics approach applied to two recent clinical isolates, CLJ1 and CLJ3, belonging to the same group of taxonomic outliers. This work gave insights into genome-wide modifications that provided bacteria with the weapons for a successful colonization and dissemination in the human host. We found that the genomes of those strains are highly dynamic and evolved within the patient due to high number of ISs. Global traits necessary for pathogenicity and survival in the host, i.e. motility, adhesion and resistance to antimicrobials, are modulated by the CLJ-ISL3 element. The later isolate CLJ3 acquired resistance toward antibiotics provided to the patients during hospitalization, with some ISs directly affecting components playing a role in resistance. Compared to the early colonizer CLJ1, the CLJ3 strain also displays higher intracellular levels of c-di-GMP, higher expression of the biofilm-associated adhesion protein CdrA, and has lost part of the LPS by the deletion of the *galU* gene; these are all features of adaptation to the so-called “chronic” lifestyle. Moreover, using bioinformatics screens we found 35 additional IS elements in the CLJ3 genome and these very likely contribute to other phenotypical changes not assessed in this study. In addition to previously known virulence determinants, numerous differentially expressed genes were annotated as “hypothetical” by using automated annotation technology RAST and their contribution in *P. aeruginosa* adaptation to human host should be further explored.

Although the exact origin of the ISL3 in the CLJ lineage is unknown, we can speculate on its acquisition from an environmental bacterial species present in the common microbial community. Indeed, the GC content of the CLJ1-ISL3 is 54.8%, while the rest of the *P. aeruginosa* genome is 66.6%, suggesting recent acquisition by a horizontal transfer. The initial disruption of the genes encoding flagellar components and pili may have allowed the CLJ strain to overcome the human immune defenses. Increased capacity to secrete the pore-forming toxin Exolysin gave the strain additional advantage to further damage the host epithelium and endothelium tissues and disseminate.

Previous studies indicated that the contribution of ISs to the adaptation of *P. aeruginosa* to CF environment is low with limited transposition events during chronic infection (93), which is in contrast to what was found in *in vitro* evolution experiment in *Escherichia coli* (94, 95). Search in the Pseudomonas data base (27) with the nucleotide sequence of the CLJ1-ISL3 fragment revealed altogether 12 strains with 100% identical fragments; 8 from Copenhagen University Hospital (96) and the others isolated from tertiary care hospital’s intensive care units in Seattle (97). The exact positions of the ISL3 element in those genomes and phenotypes related to disrupted gene(s) are currently unknown, but this knowledge would give insights on whether and how the ISL3 participated in colonization process and adaptation of those strains to a particular infectious niche. More recent genomic characterization of environmental *P. aeruginosa* isolates from dental unit waterlines, showed that in addition of the OSA loci, the IS*Pa*11 fragment altered genes of two master regulators, LasR and GacS, supporting the idea of ecological adaptive potential of *P. aeruginosa* by mobile elements (48). Therefore, in addition to small nucleotide changes in pathoadaptive genes, mobile genetic elements drive the emergence of phenotypic traits leading to adaptation of *P. aeruginosa* to niche the bacteria encounter and undoubtedly take part in strain-specific pathogenicity. The contribution of ISs to the increase of the bacterial pathogenic arsenal is likely underestimated due to still limited availability of closed bacterial genomes. New DNA sequencing technologies accessing genome fragments of tens and even hundreds of kb should reveal the global impact of mobile elements in bacterial evolution.

## Author’s contributions

ES performed all presented bioinformatics analyses of the data. PB, AB, GB, YC and AA performed experiments. SL provided materials. SE and IA designed the research and performed experiments. YC, SE and IA analyzed the data. SE and IA wrote the manuscript. All authors participated in preparing figures and tables and revised the final version of the manuscript.

## Acknowledgements

The authors thank Maria Guillermina Casabona for initial proteomic experiments on CLJ strains and Yves Vandenbrouck (CEA, Grenoble), Frédéric Boyer (UGA, Grenoble) and François Lebreton (HMS, Boston) for advice on genome analysis. Special thanks go to Mylène Robert-Genthon and Florian Chenavier for performing RT-qPCRs and PCRs and to Peter Panchev for carrying out *Galleria* infections. We are grateful to Tim Tolker-Nielsen for the generous gift of pUCP22-p*CdrA-gfp*(ASV)^c^ plasmid.

This work was supported by grants from Agence Nationale de la Recherche (ANR-15-CE11-0018-01), the Laboratory of Excellence GRAL (ANR-10-LABX-49-01) and the Fondation pour la Recherche Médicale (Team FRM 2017, DEQ20170336705). Proteomic experiments were partly supported by the Proteomics French Infrastructure (ANR-10-INBS-08-01 grant). We further acknowledge support from CNRS, INSERM, CEA, and University Grenoble Alpes. Work in SL’s laboratory was supported by a grant from the Cystic Fibrosis Foundation. AB received a Ph.D. fellowship from the CEA Irtelis program. The funding bodies have no roles in the design of the study and collection, analysis, and interpretation of data and in writing the manuscript.

**Supplementary information is available at the journal’s website**

## Supplementary Materials and Methods

### Genome sequencing, assembly, annotation and comparison

The DNA extracted from the CLJ1 bacteria was sequenced on Illumina HiSeq 2000 systems at the Beijing Genomics Institute, China with a 2×50 base-pairs (bp) paired-end mode, generating 12,333,368 reads with a genome coverage of 90-100x. PacBio (Base Clear, Leiden, Netherlands) technology was also used, providing 114,707 reads. The Illumina reads were trimmed to remove low quality sequences (limit = 0.05) and ambiguous nucleotides (maximum two nucleotides allowed) and assembled *de novo* using CLC Genomics Workbench 9.0 (Qiagen, Aarhus, Denmark). The resulting contigs were then combined with the PacBio reads using the “Join Contigs” function of the CLC Genome Finishing Module version 1.6.1 (Qiagen). Contigs or scaffolds consisting of fewer than 100 reads were filtered out and BLAST (1) searches were performed to check and remove those that show no match to *Pseudomonas* sequence in GenBank (2). CLJ3 genomic DNA was sequenced using Illumina MiSeq at the Biopolymers Facility, Harvard Medical School, Boston, USA, with 150-bp single-end runs, generating 2,495,173 reads. Quality trimming was done using CLC Genomics Workbench 9.0 using the same parameters as those used for CLJ1, followed by *de novo* assembly with minimal contig length of 1 kb. The order of both CLJ1 and CLJ3 scaffolds or contigs was determined based on the genome sequence of PA7 strain using the “move contigs” tool of the Mauve Genome Alignment Software version snapshot_2015-02-13 (3). Average nucleotide identity between genomes were estimated using ANI calculator tool on the enveomics collection server (4) with minimum alignment length, identity, and number of 700 bp, 70%, and 50, respectively, using 1 000 bp window size and 200 bp step size. SNPs between CLJ1 and CLJ3 genomes were detected by mapping trimmed CLJ3 reads to CLJ1 genome in CLC Genomics Workbench 9.0, followed by using Basic Variant Detection 1.71 tool in the Workbench with the following parameters: Ploidy = 1, Ignore positions with coverage above = 100 000, Restrict calling to target regions = Not set, Ignore broken pairs = Yes, Ignore non-specific matches = Reads, Minimum coverage = 10, Minimum count = 2, Minimum frequency (%) = 35.0, Base quality filter = Yes, Neighborhood radius = 5, Minimum central quality = 20, Minimum neighborhood quality = 15, Read direction filter = No, Relative read direction filter = Yes, Significance (%) = 1.0, Read position filter = No, Remove pyro-error variants = No. COG annotation was done using the WebMGA server (5) with E-value cutoff of 0.001. Subcellular localizations of PA7 proteins were retrieved from the Pseudomonas Genome Database, whereas those of CLJ1 and CLJ3 were predicted using PSORTb version 3.0.2 (6) and LocTree3 (7). Othologous genes among the strains were identified using OrthoMCL version 2.0.9 (8), with a BLASTp E-value cutoff of 1×10^−5^ and the default Markov cluster algorithm (MCL) inflation parameter of 1.5.

### Transcriptome analysis

The preparations of the Illumina libraries and sequencing were done by standard procedures at the Biopolymer Facility, Harvard Medical School, Boston, USA. Illumina HiSeq was used for the sequencing with 50-bp single-end runs, generating 3,225,727 and 21,633,018 reads for CLJ1 and 56,804,547 and 2,139,035 reads for CLJ3. For each replicate, raw RNA-Seq read trimmings were done in CLC Genomics Workbench 9.0 using the same parameters as for genome sequencing reads, and the trimmed reads were mapped to the annotated CLJ1 genome using the RNA-Seq analysis tool. Total number of reads mapped to the genes were incorporated into a tabular format and analyzed using the DESeq2 differential expression analysis pipeline (9). Differentially expressed genes between CLJ1 and CLJ3 were identified using a 5% False Discovery Rate (FDR).

### Proteomics

<u>Sample preparation.</u> Overnight cultures of CLJ1 were diluted to an OD600 0.1 in 30 mL, CLJ3 was left a night at room temperature and then the cultures were incubated at 37°C under shaking to an OD600 0.8. At this point, 30 mL of bacterial cultures were centrifuged at 6,000 rpm, 4°C for 10 min, and supernatants were filtered with 0.22 μm filters. Total <u>membranes separation.</u> Pellets were re-suspended in 1 mL of 10 mM Tris-HCl, 20 % sucrose, pH8 buffer supplemented with protease inhibitors cocktail (PIC, Roche, Basel, Switzerland) and were disrupted by sonication. Unbroken bacteria were removed by a centrifugation at 8,000 rpm for 10 min at 4°C. Total membrane fraction was obtained by ultracentrifugation at 200,000 g for 1 h at 4°C, and the pelleted fraction was washed twice with 1 mL of 10 mM Tris-HCl, 20 mM MgCl2, pH8, supplemented with PIC, and resuspended in 500 μL of the same buffer. <u>Supernatant fraction.</u> Proteins from supernatants were precipitated by a TCA-sarkosyl method (0.5% final volume of sarkosyl and 7.5% final volume of TCA) for 2 h on ice and centrifuged at 12,000 rpm for 15 min. Pellets were washed twice with tetrahydrofuran, and re-suspended in 50 μL of loading buffer. Prepared samples, in triplicates, were then analyzed by SDS-PAGE and immunoblotting using antibodies directed against 3 synthetic peptides of ExlA ((10), 1:1 000), anti-RpoA (1:5000) as control for whole cell, anti-TagQ ((11), 1:10 000) as control for the membrane fraction and anti-DsbA (1:2 000) as control for the periplasm fraction. Secondary antibodies used were HRP-coupled anti-rabbit (1:50 000) and anti-mouse (1:5 000) (Sigma-Aldrich). Western blots were developed using Luminata Crescendo Western HRP (Millipore) substrate.

<u>Mass spectrometry-based quantitative proteomic analyses.</u> Extracted proteins were prepared as described in (12). Briefly, proteins were stacked in the top of a SDS-PAGE gel (NuPAGE 4-12%, ThermoFisher Scientific), stained with Coomassie blue (R250, Bio-Rad) before in-gel digestion using modified trypsin (Promega, sequencing grade). Resulting peptides were analysed by nanoliquid chromatography coupled to tandem mass spectrometry (Ultimate 3000 coupled to LTQ-Orbitrap Velos Pro, Thermo Scientific) using a 120-min gradient (2 analytical replicates per biological replicate). RAW files were processed using MaxQuant (13) version 1.5.3.30. The protein content in total, membrane, and secretome proteomes of CLJ1 and CLJ3 were analyzed independently from the others. Spectra were searched against the homemade CLJ database and the frequently observed contaminants database embedded in MaxQuant. Trypsin was chosen as the enzyme and 2 missed cleavages were allowed. Peptide modifications allowed during the search were: carbamidomethylation (C, fixed), acetyl (Protein N-ter, variable) and oxidation (M, variable). Minimum peptide length was set to 7 amino acids. Minimum number of peptides, razor + unique peptides and unique peptides were all set to 1. Maximum false discovery rates - calculated by employing a reverse database strategy ‐were set to 0.01 at peptide and protein levels. Intensity-based absolute quantification values iBAQ (14) were calculated from MS intensities of unique+razor peptides. Statistical analyses were performed using ProStaR (15). Proteins identified in the reverse and contaminant databases, proteins only identified by site and proteins exhibiting less than 3 iBAQ values in one condition were discarded from the list. After log2 transformation, intensity values were normalized by median centering before missing value imputation (replacing missing values by the 2.5 percentile value of each column); statistical testing was conducted using *limma* t-test. Differentially recovered proteins were sorted out using a log2(fold change) cut-off of 2 and a FDR threshold on remaining p-values of 1% using the Benjamini-Hochberg procedure. The different lists were then combined together. In total, this allows us to end up with a list of 2 852 quantified proteins.

### RT-qPCR

Yield, purity and integrity of RNA were evaluated on Nanodrop and by agarose gel migration. Complementary DNA synthesis was carried using 3 μg of RNA with SuperScript III First-Strand Synthesis System (Invitrogen) with or without SuperScript III RT enzyme to assess the absence of genomic DNA. The CFX96 Real-Time system (BioRad) was used to amplify the cDNA and the quantification was based on use of SYBR green fluorescent molecules. cDNA was incubated with 5 μL of Gotaq qPCR master mix (Promega) and reverse and forward specific primers at a final concentration of 125 nM in a total volume of 10 μL. Cycling parameters of the real time PCR were 95°C for 2 min, 40 cycles of 95°C for 15 s and 60°C for 45 s, and finally a melting curve from 65°C to 95°C by increment of 0.5°C for 5 s to assess the specificity of the amplification. To generate standard curves, serial dilutions of cDNA pool of the CLJ strains were used. The experiments were performed with three biological samples for each strain, in duplicate, and the results were analyzed with the CFX manager software (BioRad). The relative expression of mRNAs was calculated using the DDCq method relative to *rpoD* reference Cq values. The graph was represented with the SEM (Standard Error of the Mean). Statistical analysis was carried out using a non-parametric Mann-Whitney U test. The *p*values < 0.05 were considered as significant.

---

**Fig. S1.**
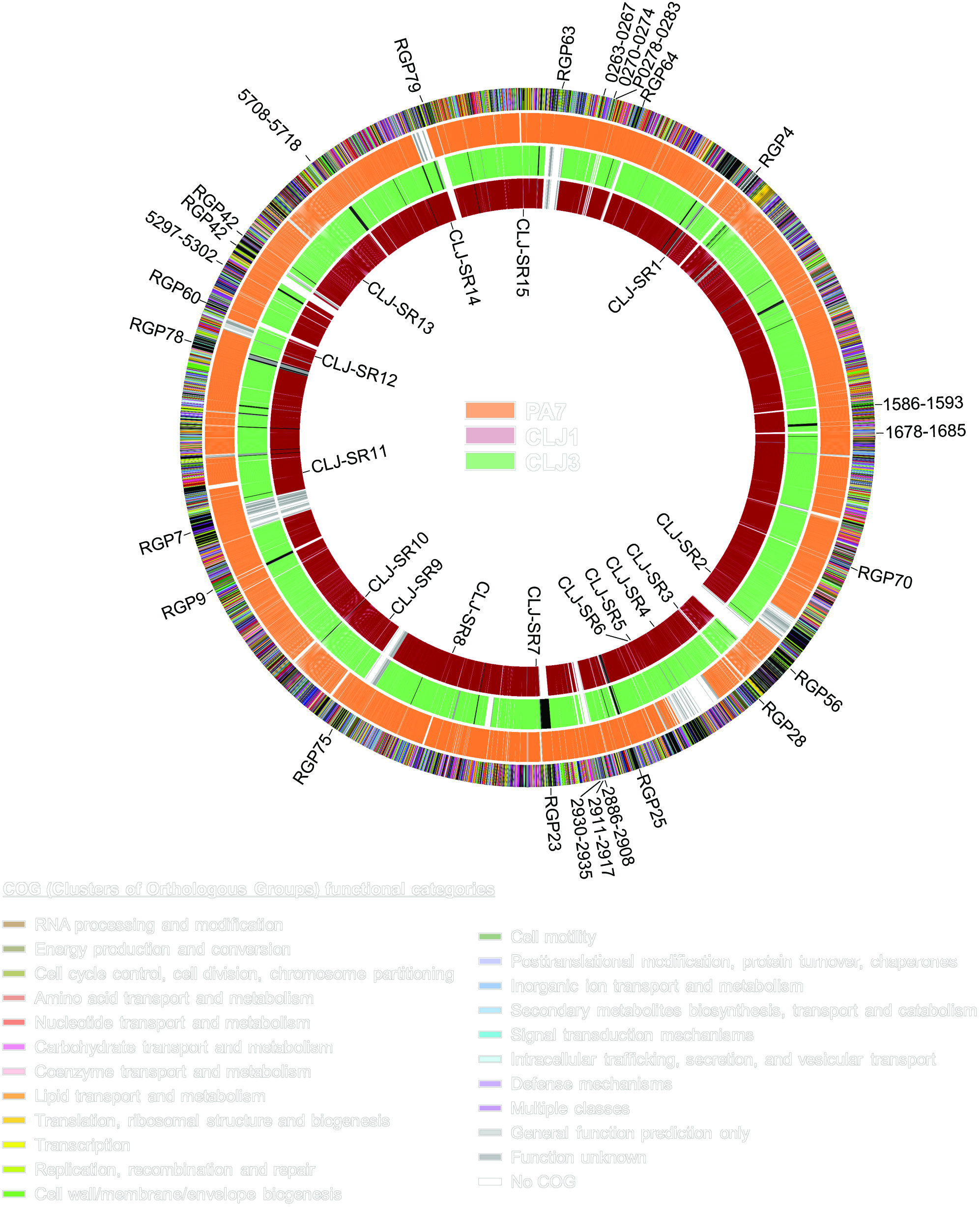
**Details of whole genome comparison between PA7 and CLJ strains.** The outer ring shows all the genes in the strains colored according to their COG (Clusters of Orthologous Groups) functional categories as listed on the bottom, and the three other rings represent the PA7 (orange), CLJ3 (green), and CLJ1 (red) genomes, respectively. White color indicates that the gene is absent from the genome, grey color indicates that the gene or its homolog is present in a different position in the genome, while black color in the CLJ rings represents gaps between contigs. The inner labels show the CLJ-specific regions and the outer labels are the PA7-specific regions (represented in RGPs or PSPA7 locus numbers) described in Additional file 2: Table S1 and Table S2, respectively. (PDF 6909 KB)

**Table S1.** Genome assembly and annotation statistics (DOCX 17Ko)

|  | CLJ1 | CLJ3 |
| --- | --- | --- |
| **Number of contigs/scaffolds** | 68 | 135 |
| **Average coverage (×)** | 98.3 | 96.9 |
| **Contig/scaffold size (bp)** | 201-847201 | 1025-297138 |
| **Average contig/scaffold size (bp)** | 95801 | 47065 |
| **Contig/scaffold N50 (bp)** | 456227 | 97517 |
| **Total genome size (bp)** | 6514448 | 6353726 |
| **G+C content (%)** | 66.6 | 66.6 |
| **Protein coding sequences** | 6259 | 6107 |
| **tRNA** | 62 | 57 |
| **rRNA** | 9 | 3 |

**Table S2.**
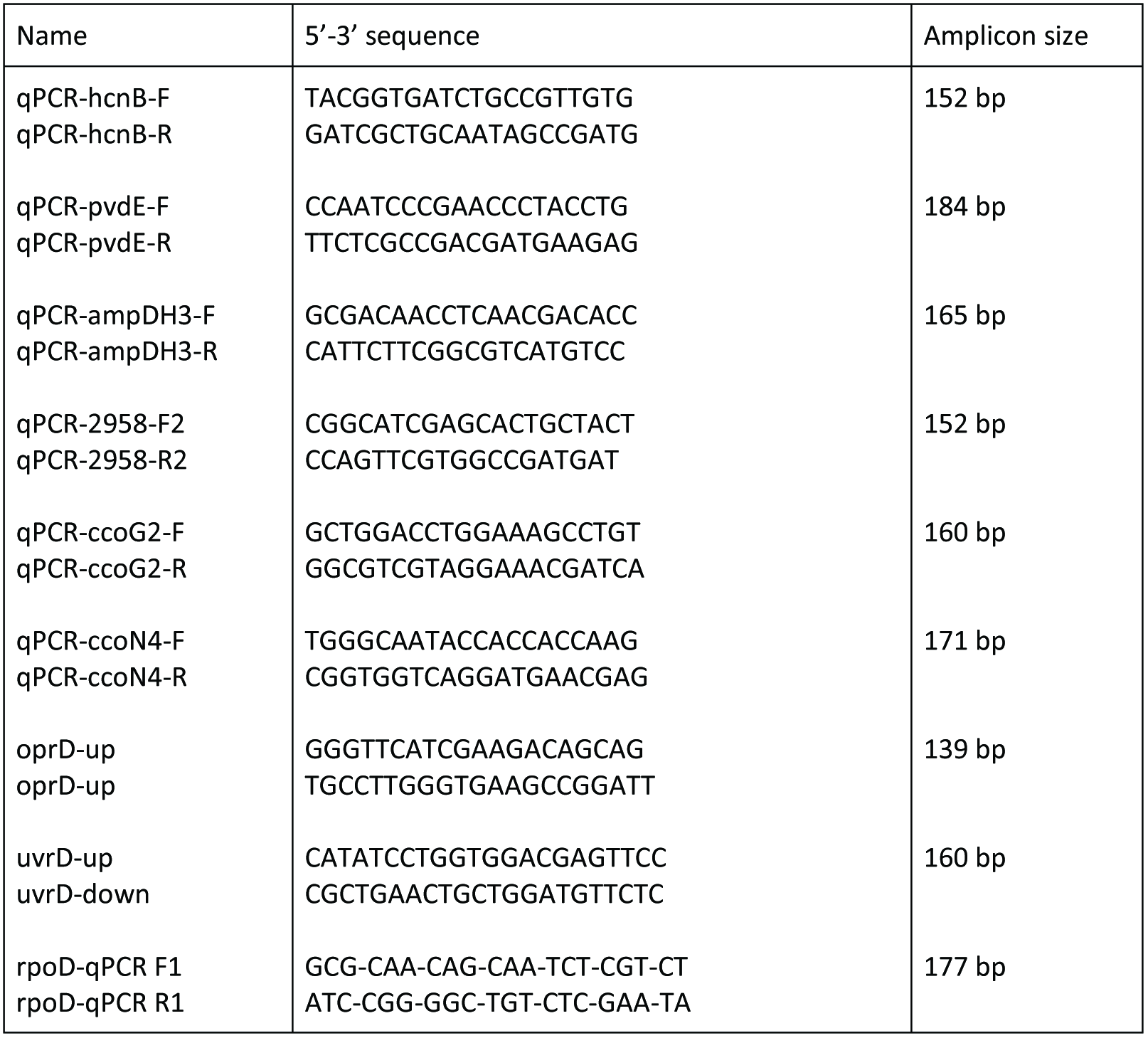
RT-qPCR primers used in this study (DOCX 16Ko)

**Table S3.** Specific regions of CLJ compared to PA7 (DOCX 17Ko)

| Region | RGP | CLJ1 locus | Number of genes | Features |
| --- | --- | --- | --- | --- |
| CLJ-SR1 | RGP66 | 0487-0494 | 8 | Phage-related |
| CLJ-SR2 | RGP29* | 2128-2214 | 87 | PAGI-2-like island |
| CLJ-SR3 | RGP28 | 2316-2322 | 6 | Mobile element proteins |
| CLJ-SR4 | RGP27 | 2415-2573 | 160 | Dit island |
| CLJ-SR5 | RGP72 | 2579-2583 | 5 | D-galactonate catabolism |
| CLJ-SR6 | RGP26 | 2615-2621 | 7 | Phage-related; lytic enzymes |
| CLJ-SR7 | RGP23 | 2917-2924 | 7 | Mobile element proteins; possible DNA helicase |
| CLJ-SR8 |  | 3303-3309 | 7 | ABC transporter ATP-binding protein |
| CLJ-SR9 |  | 3607-3616 | 9 |  |
| CLJ-SR10 | RGP15 | 3774-3784 | 11 | Threonine dehydratase; ABC transporter proteins |
| CLJ-SR11 |  | 4314-4301 | 14 | AlpBCDE-lysis cassette, phage-related |
| CLJ-SR12 |  | 4838-4876 | 39 | Pyrroloquinoline quinone synthesis proteins |
| CLJ-SR13 |  | 5192-5203 | 10 | Phage related; accessory cholera enterotoxin |
| CLJ-SR14 |  | 5811-5871 | 55 | Heavy metal resistance |
| CLJ-SR15 |  | 6156-6161 | 6 | Phage-related |

RGP: region of genomic plasticity, \*CLJ strains do not share any genes with PA7 in the RGP

**Table S4.**
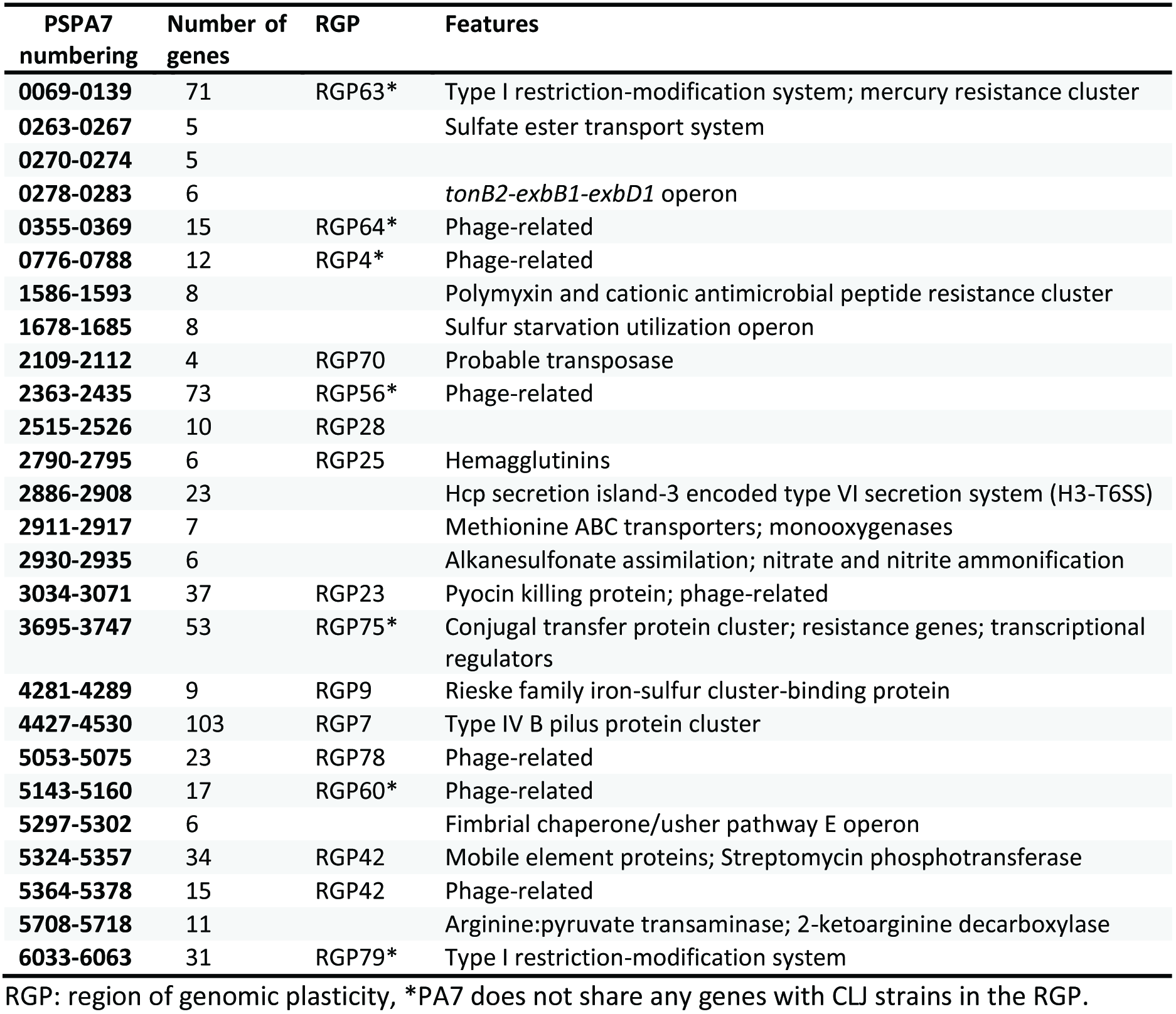
Specific regions of PA7 compared to CLJ (DOCX 18Ko)

**Table S5.** Genes in CLJ-SR14 (DOCX 18Ko)

| PA7 | CLJ1 | CLJ3 | RAST annotation |
| --- | --- | --- | --- |
| NA | CLJ1_5811 | CLJ3_5608 | MG(2+) CHELATASE FAMILY PROTEIN / ComM-related protein |
| NA | CLJ1_5814 | CLJ3_5609 | hypothetical protein |
| NA | CLJ1_5815 | CLJ3_5610 | hypothetical protein |
| NA | CLJ1_5817 | CLJ3_0003 | hypothetical protein |
| NA | CLJ1_5818 | CLJ3_0004 | Error-prone, lesion bypass DNA polymerase V (UmuC) |
| ns | CLJ1_5819 | CLJ3_0005 | hypothetical protein |
| ns | CLJ1_5820 | CLJ3_0006 | Mercuric ion reductase (EC 1.16.1.1) |
| ns | CLJ1_5821 | CLJ3_0007 | Mercuric transport protein, MerC |
| ns | CLJ1_5822 | CLJ3_0008 | Periplasmic mercury(+2) binding protein |
| ns | CLJ1_5823 | CLJ3_0009 | Mercuric transport protein, MerT |
| ns | CLJ1_5824 | CLJ3_0010 | Mercuric resistance operon regulatory protein |
| ns | CLJ1_5825 | CLJ3_0011 | hypothetical protein |
| NA | CLJ1_5826 | CLJ3_0012 | Sterol desaturase |
| NA | CLJ1_5827 | CLJ3_0013 | Transcriptional regulator, AraC family |
| NA | CLJ1_5828 | CLJ3_0014 | Lipoprotein signal peptidase (EC 3.4.23.36) |
| NA | CLJ1_5829 | CLJ3_0015 | hypothetical protein |
| NA | CLJ1_5830 | CLJ3_0016 | Cobalt-zinc-cadmium resistance protein CzcD |
| NA | CLJ1_5831 | CLJ3_0017 | COG3267: Type II secretory pathway, component ExeA (predicted ATPase) |
| NA | CLJ1_5832 | CLJ3_0018 | FIG131328: Predicted ATP-dependent endonuclease of the OLD family |
| ns | CLJ1_5833 | CLJ3_0019 | Mobile element protein |
| ns | CLJ1_5834 | CLJ3_0020 | Mobile element protein |
| NA | CLJ1_5836 | CLJ3_6032 | transcriptional regulator MvaT, P16 subunit, putative |
| ns | CLJ1_5837 | CLJ3_6031 | Gifsy-2 prophage protein |
| ns | CLJ1_5838 | CLJ3_6030 | Error-prone repair protein UmuD |
| ns | CLJ1_5839 | CLJ3_6029 | Error-prone, lesion bypass DNA polymerase V (UmuC) |
| NA | CLJ1_5840 | CLJ3_6028 | putative (L31491) ORF2; putative [Plasmid pTOM9] |
| NA | CLJ1_5841 | CLJ3_6027 | putative ORF1 [Plasmid pTOM9] |
| NA | CLJ1_5842 | CLJ3_6026 | NreA-like protein |
| NA | CLJ1_5843 | CLJ3_6025 | Inner membrane protein |
| NA | CLJ1_5844 | CLJ3_6024 | probable membrane protein YPO3302 |
| NA | CLJ1_5845 | CLJ3_6023 | hypothetical protein |
| NA | CLJ1_5846 | CLJ3_6022 | hypothetical protein |
| NA | CLJ1_5847 | CLJ3_6021 | Chromate transport protein ChrA |
| NA | CLJ1_5848 | CLJ3_6020 | Chromate resistance protein ChrB |
| NA | CLJ1_5849 | CLJ3_6019 | Phage integrase family protein |
| NA | CLJ1_5851 | CLJ3_6018 | RuBisCO operon transcriptional regulator CbbR |
| NA | CLJ1_5852 | CLJ3_6017 | Phosphonate dehydrogenase (EC 1.20.1.1) (NAD-dependent phosphite dehydrogenase) |
| NA | CLJ1_5853 | CLJ3_6016 | Phosphonate ABC transporter permease protein phnE (TC 3.A.1.9.1) |
| NA | CLJ1_5854 | CLJ3_6015 | Phosphonate ABC transporter phosphate-binding periplasmic component (TC 3.A.1.9.1) |
| NA | CLJ1_5855 | CLJ3_6014 | Phosphonate ABC transporter ATP-binding protein (TC 3.A.1.9.1) |
| ns | CLJ1_5856 | CLJ3_6013 | FIG002188: hypothetical protein |
| ns | CLJ1_5857 | CLJ3_6012 | FIG067310: hypothetical protein |
| NA | CLJ1_5858 | CLJ3_6011 | hypothetical protein |
| ns | CLJ1_5859 | CLJ3_6010 | Long-chain-fatty-acid--CoA ligase (EC 6.2.1.3) |
| NA | CLJ1_5860 | gap | Enoyl-CoA hydratase (EC 4.2.1.17) |
| NA | CLJ1_5861 | ns | Mobile element protein |
| NA | CLJ1_5862 | ns | Mobile element protein |
| NA | CLJ1_5863 | gap | transcriptional regulator, TetR family |
| NA | CLJ1_5864 | CLJ3_5612 | Error-prone, lesion bypass DNA polymerase V (UmuC) |
| NA | CLJ1_5866 | ns | Mobile element protein |
| NA | CLJ1_5867 | CLJ3_5613 | hypothetical protein |
| NA | CLJ1_5868 | CLJ3_5614 | Mobile element protein |
| NA | CLJ1_5869 | CLJ3_5615 | hypothetical protein |
| NA | CLJ1_5870 | CLJ3_5616 | hypothetical protein |
| NA | CLJ1_5871 | CLJ3_5617 | hypothetical protein |

NA: no orthologous gene is found in the corresponding genome, ns: orthologous gene(s) is/are found in other location(s) in the corresponding genome, gap: the gene location corresponds to a gap between contigs in the corresponding genome.

**Table S6.**
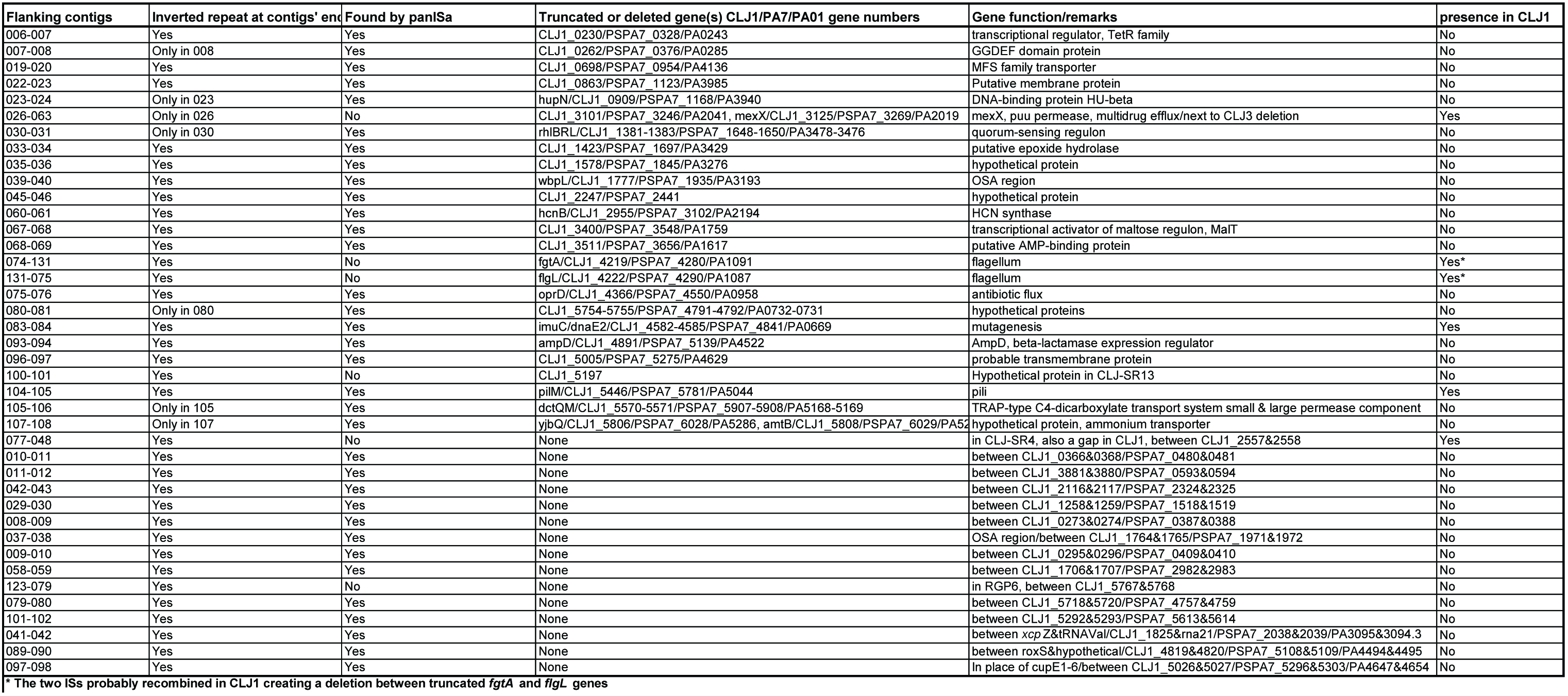
Prediction of CLJ-ISL3 insertions between contigs of CLJ3 (XLSX 14Ko)

**Table S7** Differential proteomic analysis between CLJ1 and CLJ3 in total proteome, membrane proteome, and secretome proteome (XLSX 376Ko)

**Table S8** Differential gene expression analysis between CLJ1 and CLJ3 (XLSX 436Ko)

**Table S9** List of genes/proteins that are statistically significantly differentially expressed between CLJ1 and CLJ3 in both, RNA-Seq and in at least one of the proteomic datasets (XLSX 24K0)

**Table S10.** CLJ phage-related region (DOCX 16Ko)

| CLJ1 locus | Region | Number of genes | Corresponding PA7 locus | Mobility gene present |
| --- | --- | --- | --- | --- |
| **0462-0497** | RGP66 (including CLJ-SR1) | 36 | PSPA7_0678-0716 | Integrase |
| **0535-0557** | RGP3 | 23 | PSPA7_0754-0775 | None |
| **2607-2624** | RGP26 (including CLJ-SR6) | 18 | PSPA7_2648-2661 | Integrase |
| **4295-4314** | Including CLJ-SR11 | 20 | PSPA7_4602-4606 | None |
| **4778-4791** | RGP78 | 14 | PSPA7_5040-5080 | Integrase |
| **5192-5203** | CLJ-SR13 | 12 | NA | Integrase |
| **5758-5774** | Including RGP6 | 17 | PSPA7_4699-4703 | Integrase |
| **6156-6161** | CLJ-SR15 | 6 | NA | None |

NA: not applicable

